# *Afrothismiaceae* West of the Dahomey gap: *Afrothismia fonensis* sp. nov. Critically Endangered and endemic to Pic de Fon forest, Simandou, Republic of Guinea

**DOI:** 10.64898/2026.05.05.723002

**Authors:** Martin Cheek, Denise Molmou, Guillaume Delhaye

## Abstract

The fully mycoheterotrophic, non-photosynthetic *Afrothismia fonensis* Cheek & G.Delhaye sp. nov. (Afrothismiaceae), is described and illustrated from two sites in submontane forest in or adjacent to the Pic de Fon Forêt Classée, Simandou Range, Republic of Guinea. This is the first record of the genus and family in West Africa west of Nigeria. The new species is remarkable for its small size, and for being unique in the genus in the entirely connate intertepaline lobes (in other species of the genus they are free or only partly united) and the longitudinal ridges on the outer perianth tube (unknown in other species). The provisional extinction risk assessment for *Afrothismia fonensis* is Critically Endangered (CR B1ab (iii)+2ab(iii)+D1) using the IUCN 2012 categories and criteria, due to less than 50 individuals being recorded, and due to the both the very small range and the immediate threats from foraging by red river hogs, trampling by cattle and from de-watering of the adjacent Oueleba iron-ore body where mining began in 2025. It should be noted that mitigation actions are expected to adequately address the risks associated with mining activities, and direct impacts to both areas of *Afrothismia fonensis* habitat have been fully avoided through relocation of planned infrastructure.

We review the importance of the Boyboyba forest, Simandou range, as the West African centre of diversity for non-photosynthetic heteromycotrophs. This new discovery is examined in the context of other recently discovered range extensions to Guinea of Central African genera and families.

## Introduction

Fully mycoheterotrophic non-photosynthetic plants, also known as achlorophyllous mycotrophic plants, mycoheterophs (Merckx 2013) or saprophytes, are remarkable for lacking chlorophyll and being completely dependent on fungi for their supply of carbon. In continental Africa, individual species or entire genera that fit this description occur in the families Burmanniaceae, Gentianaceae and Orchidaceae, while all members of Afrothismiaceae, Thismiaceae and Triuridaceae are fully mycotrophic (Cheek & Ndam 1996; Cheek & Williams 1999; Cheek 2006; Cheek et al. 2023).

Although some earlier authors (Jonker 1938; Maas-van der Kamer 1998) placed *Afrothismia* (Engl.) Schltr. and associated genera as a tribe Thismieae within the family Burmanniaceae sensu lato, molecular phylogenetic data (e.g. Merckx et al. 2006) strongly indicated that Thismiaceae are best placed as a separate family (Cheek et al. 2018a). Subsequently, well-sampled analyses placed *Afrothismia* as sister to Taccaceae + Thismiaceae, in a different subclade of Dioscoreales from Burmanniaceae *sensu stricto* (Merckx et al. 2009; Lin et al. 2022). These groups are separated from each other by numerous morphological characters and the family Afrothismiaceae was formally recognised (Cheek et al. 2023).

*Afrothismia* are confined to the humid forests of tropical continental Africa. *Afrothismia* differs from the genera of Thismiaceae by the annulus inserted deep inside the perianth tube; stamens inserted below the annulus; anthers usually adnate to stigma (but note that anthers adnate to stigmas have recently been reported in the new Thismiaceae genus *Relictithismia* Suetsugu & Tagane (Suetsugu et al. 2024)); and rhizomes with clusters of ± spherical tubers, among other characters (Cheek et al. 2023). Seventeen species have been formally described (with this paper 18), but there appear to be seven additional undescribed species mainly in Gabon and Cameroon (Cheek et al. 2023).

The genus *Afrothismia* was erected by Schlechter (1906), based on *Thismia winkleri* Engl. (Engler 1905), with *A. pachyantha* Schltr., both from Cameroon. The range of the genus was formally extended to East Africa by Cowley (1988), with *A. insignis* Cowley from Tanzania and *Afrothismia winkleri* var. *budongensis* Cowley (now *A. ugandensis* Cheek (Cheek et al. 2024a) from Uganda. Publication of the Cameroonian species, *A. gesnerioides* H.Maas (Maas-van der Kamer 2003) doubled the number of the original two species known for the genus from nearly a century earlier, to four. In the next 16 years, the number of formally described new species quadrupled to 16, with discoveries from Gabon (Dauby et al. 2008),

Kenya (Cheek 2004), Malawi (Cheek 2009) and Tanzania (Cheek & Jannerup 2006); but the largest number of discoveries by far has been made in Cameroon, where eight new species of *Afrothismia* have been published, one extending to Nigeria (Franke 2004; Franke et al. 2004; Sainge & Franke 2005; Sainge et al. 2005; Cheek 2007; Sainge et al. 2013; Cheek et al. 2019). Most of the Cameroonian species fall within the Cross-Sanaga interval (Cheek *et al*. 2001), which holds the highest flowering plant species (Barthlott et al. 1996) and generic (Dagallier et al. 2020) diversity per degree square in Tropical Africa, and which has been the site of many new discoveries, including genera new to science (Litt & Cheek 2002; Cheek et al. 2003; Cheek et al. 2018b). Eight of the Cameroonian *Afrothismia* species feature in the Red Data Book of Cameroon Plants (Onana & Cheek 2011) and most species of *Afrothismia* are Critically Endangered due to their extremely restricted distributions (Cheek et al. 2023). One Cameroonian species is considered extinct after targeted searches (Cheek et al. 2019), while several others have not been recorded alive in several decades and so may also be extinct, e.g. *A. zambesiaca* Cheek (Cheek 2009).

Although species niche modelling had indicated that *Afrothismia* might occur in West Africa, west of the Dahomey gap, the ‘highly suitable’ areas predicted were not in Guinea but in Liberia, Cote D’Ivoire and Sierra Leone (Sainge et al. 2017). Previous targeted expert surveys for *Afrothismia* in Guinea failed, although they produced other new species of fully mycotrophic Dioscoreales (Cheek & van der Burgt 2010; Cheek et al. 2024b).

Lacking any green tissue, *Afrothismia* spp. depend on vascular arbuscular mycorrhizal fungi for sustenance. The genus *Rhizophagus* P.A. Dang (Glomerales, Glomeromycota) has been implicated as the fungal partner of the genus (Franke et al. 2006). Delayed co-speciation between *Afrothismia* and the fungal partner has been demonstrated (Merckx 2008; Merckx & Bidartondo 2008). The autotrophic partners of these fungi remain unknown but research on *Thismia* suggests that mycoheterotrophic plant could target specific lineages of mycorrhizal fungi compared to neighbouring green plants (Gomes et al. 2017).

The new species of *Afrothismia* reported in this paper is a result of a field study by the three authors which was initiated in October 2025 to find the mycorrhizal and autotrophic partners of the Critically Endangered *Gymnosiphon fonensis* Cheek (Cheek et al. 2024b) known globally from five sites in the Pic de Fon Forêt Classée (PDF FC) to which it appears endemic. The new *Afrothismia* was found growing with it at two of these sites, first by GD at the ‘tunnel forest’ and then at Boyboyba forest by DM.

## Materials & methods

Nomenclature follows the *Madrid Code* (Turland et al. 2025). Names of species and authors follow IPNI (continuously updated). Herbarium material was examined with a Leica Wild M8 dissecting binocular microscope fitted with an eyepiece graticule measuring in units of 0.025 mm at maximum magnification. The drawing was made with the same equipment with a Leica 308700 camera lucida attachment. The following herbaria were inspected for specimens: B, BM, EA, HNG, K, MHU, SRGH, YA and WAG. Herbarium codes follow Index Herbariorum (Thiers, continuously updated).

The use of technical terms follows Beentje & Cheek (2003) and the format of the description follows those in other papers describing new species in *Afrothismia*, e.g. Cheek & Jannerup (2006), Cheek et al. (2024b). The specimens cited have all been seen by the co-authors. The conservation assessment follows the IUCN (2012) Red List of Threatened Species categories and criteria using current guidance (IUCN Standards and Petitions Committee 2024).

### Results - Taxonomy

#### *Afrothismia fonensis* Cheek & G.Delhaye sp nov. Type

Republic of Guinea, Guinée-Forestière, Simandou Range, Forêt Classée de Pic de Fon, Boyboyba Forest, Oueleba (also known as Ouéléba) to Wataferedou path, 8°40′ 14”N, 8°52′ 46.9”<u>W</u>, c. 800 m, fl.fr. 9 Oct. 2025, *Cheek* with Delhaye and Molmou 19408 (holo-K[K000593395]; iso-HNG).

#### Additional collection

Republic of Guinea, Guinée-Forestière, Simandou Range, to northern boundary of Forêt Classée de Pic de Fon, Tunnel Forest, 8°43′ 21”N, 8°52′ 33.4”<u>W</u>, c. 800 m, fl.fr. 8 Oct. 2025, *Cheek* with Delhaye and Molmou 19405B (HNG, K).

#### Diagnosis

differing from all other known species of *Afrothismia* Schltr. in the presence of a reflexed single intertepaline lobe between each pair of adjacent tepals (not with each tepal having a pair of hastate lobes at the base, or lacking any basal lobes) and in the presence of longitudinal ridges on the ventral side and flanks of the perianth tube (not smooth, or verrucate, or tuberculate, or otherwise ornamented).

*Achlorophyllous mycotrophic perennial herb* with the flower and fruit emerging above the leaf-litter, clusters of bulbils also usually visible. *Stem* (rhizome), succulent, translucent-white, concealed in substrate close to surface, horizontal, little branched, terete, at least 10 mm long (and possibly 10 cm or more), 0.8-0.9 mm diam., internodes 3-8 mm long. Scale-leaves translucent-white, slightly spreading, ovate-lanceolate, c. 2 × 1 mm; axillary buds globose. *Bulbil clusters* scattered along stems and inflorescence axes, isodiametric or broader than long, 2-3(−5) × 3-4(−5) mm, sometimes closely spaced, 8-10 bulbils per cluster, bulbils ellipsoid, rarely globose, each (0.5-)0.8-2 × 0.5-0.9 mm, with an apical rootlet (1.5-)6-27 mm long (Fig. 1F, Fig. 2A&C). Inflorescence terminal, cincinnal, 1-2(−4)-flowered, rhachis internodes 6-8 mm long; bracts as the scale leaves; terminal internodes (pseudo-pedicels) 1-2 mm long at anthesis, 2.5(−3) mm long in fruit. *Flowers* developing in succession as fruits develop, one flower open at a time per stem. Subtending floral bract translucent-white, enveloping the ovary and the dorsal surface of the perianth tube, extending to the upper perianth tube or as far as the base of the dorsal tepals, concave, lanceolate, 4.5–8 × 1.9–2.5 mm, apex acute. *Perianth tube* angled, mainly directed horizontally or above the horizontal, 4.2–7.2 mm long from ovary to perianth mouth, constricted slightly between lower (proximal) and (dorsal surface) upper (distal) parts of the tube; outer surface with 14-16 interrupted longitudinal ridges on ventral surface and sides, mainly on the lower perianth tube (Fig.1 A&B, 2A&C), surface otherwise smooth, glabrous. *Lower perianth tube* horizontal, subellipsoid, 4–4.2 mm long, c. 2.8 mm wide at base, 3-3.2 mm wide at midpoint, 2.5 mm wide at junction with upper perianth tube, translucent-white (sometimes with 6 longitudinal faint pink lines on the proximal part of the lower tube corresponding to the adnate filaments on the inner surface). *Annulus* inserted c. 1 mm above the free staminal filament insertion on the dorsal surface, asymmetric, unlobed, glabrous and discontinuous, absent in the ventral 1/3 of the tube; in the dorsal 2/3 of the tube projecting at 90º into the tube cavity for up to 0.1-0.15 mm, apex rounded. *Upper perianth tube* uniformly purple-red, erect, forming an angle of 90º with the dorsal surface of the lower perianth tube, shortly cylindrical, c. 3.2 × 2.5 mm, apex rounded, the mouth horizontal, angled at 90º from the axis of the upper tube, surrounded by a corona, inner and outer surfaces glabrous. Mouth orbicular, 1.5-1.7 mm diam., hemi-orbicular, or transversely elliptic-oblong. Corona 0.25-0.3 mm long, projecting forward as a rim from the insertion of the tepals around the mouth (when mouth orbicular), or partly occluding the opening of the perianth tube (when mouth elliptic-oblong). *Tepals* (5-)6, dimorphic (unequal), spreading to ± forward-directed, translucent-white, narrowly triangular-linear, 0.6-0.8(−1) mm wide at base, margins revolute at base, apex rounded, glabrous; dorsal tepal or tepal pair longest, 11-12 mm long; lateral pair 4.5-5.5 mm long; ventral tepal pair 4-5.5. mm long. Intertepal lobes (see discussion) closely resembling the tepals, but reflexed, membranous, completely translucent, margins not revolute, 1.8-2.2 × 0.4-0.6 mm, ventrally appressed to and extending to the lower perianth tube; distally slightly spreading, not adpressed. *Stamens* 6, staminal filaments joined proximally to perianth tube for 0.8-1.2 mm, sometimes appearing as dark lines on exterior tube surface, distally free, patent, red, ±terete 0.75-1.25 mm long, patent and arching inward to stigma (Fig. 1C, 2D), tapering from 0.25 mm wide at base to 0.15 mm wide distally, swollen at apex; anthers angled from filaments towards stigma, oblong-elliptic in plan view, 0.4-0.5 × 0.3 mm, in side view swollen at base, curving up and tapering towards apex (resembling a duck’s head); thecae 2, each 0.15 mm wide, adjacent, concealed by connective dorsally (Fig. 1D) but not separated by connective ventrally (Fig. 1E); distal connective appendage triangular, 0.15 × 0.15 mm, joined to undersurface of stigma (Fig. 1E). *Ovary* subturbinate, 2 × 2.5 mm; placentation axile (placenta attached at base and apex), the ovules and placental mass globose, 1.25 × 1.8 mm, inserted in the centre of the ovary. Style cylindrical, c.0.9 mm long, 0.45 mm diam., stigma discoid, inconspicuously 6-lobed, c.1 mm diam., surface papillate, glabrous.

**FIG 1.**
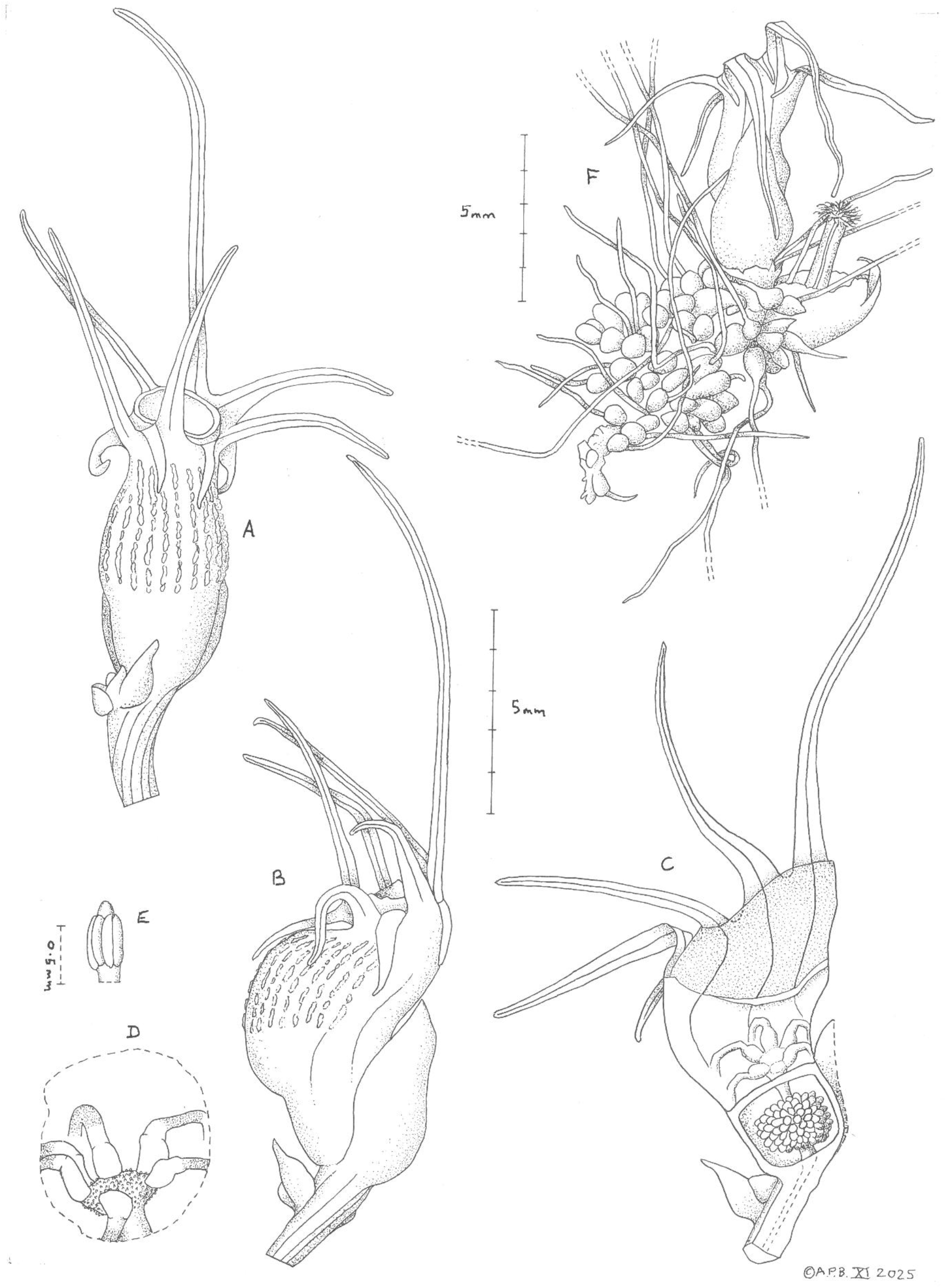
Afrothismia fonensis. A flower and stem, ventral view; B flower, side view; C longitudinal section of flower, the shaded portion = the upper perianth tube (red); D stamens attached to stigma (viewed through aperture in perianth tube); E anther, ventral surface; F view of whole plant, showing bulbils, flower in dorsal view and (right) dehisced fruit with placenta raised on ‘placentophore’. All drawn from *Cheek* with Delhaye and Molmou 19405B (spirit, K) by ANDREW BROWN.

**FIG 2.**
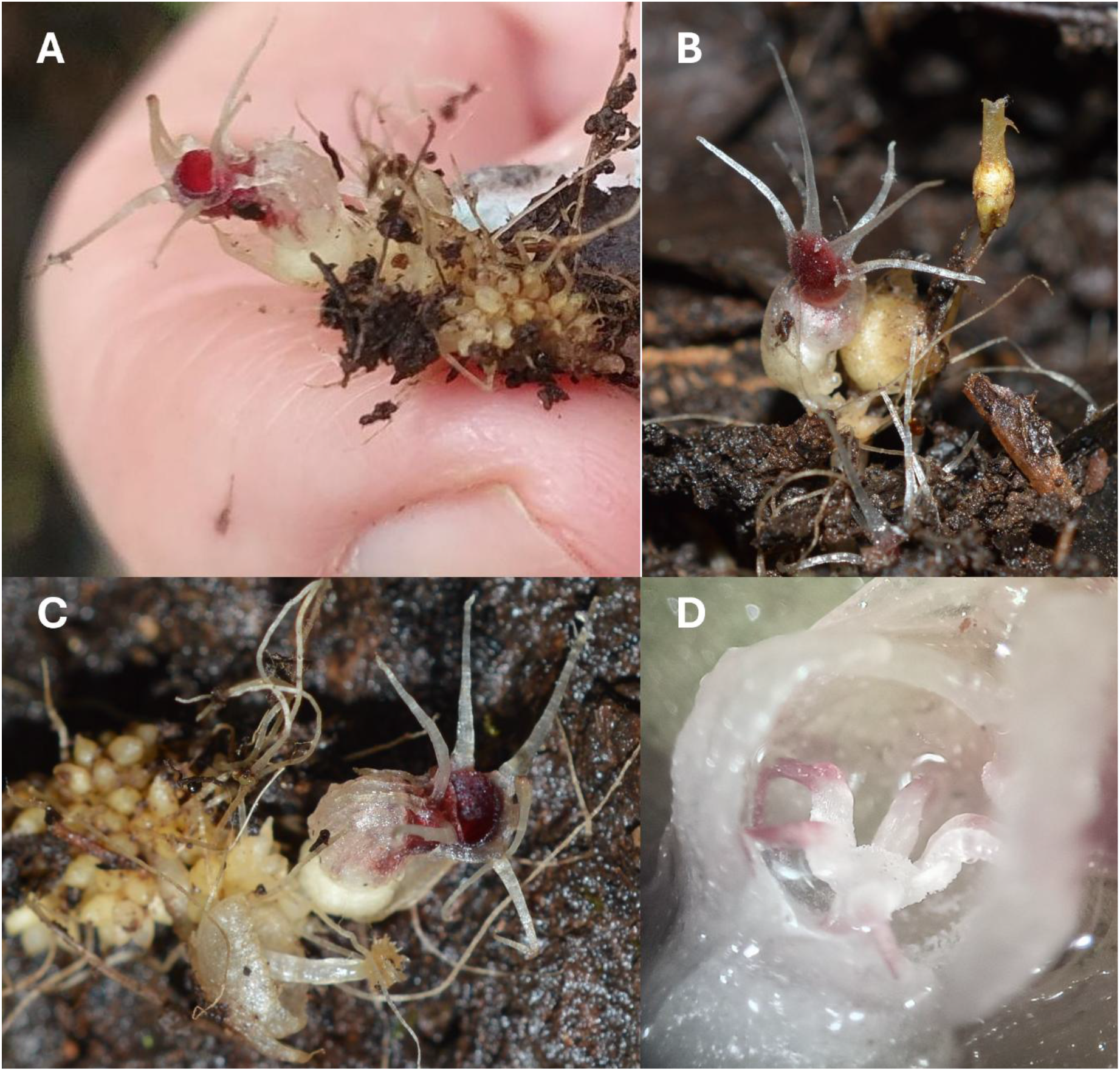
Afrothismia fonensis. A whole plant, note longitudinal ridges of perianth tube; B plant *in situ* in Boyboyba forest, dorsal view of flower, undehisced globose fruit, and (to right) *Gymnosiphon samoritoureanus*; C whole plant, note intertepaline lobes and circumscissile dehisced fruit with placentophore-elevated placental brush; D anthers, on arched free filaments, apices attached to the stigma. A-C from *Cheek* 19405B (HNG, K) Tunnel Forest, D from *Cheek* 19408 (HNG, K) Boyboyba Forest. Photos by Guillaume Delhaye (B-D) & Martin Cheek (A).

*Fruit* globose, c.4 × 3.5-4.5 mm (Fig. 2B). Circumscissile dehiscence, the lid of fruit (perianth floor) semi-orbicular, detaching from the rest of the fruit, the placental column elongating from 0.25-0.5 mm in the ovary to 4-5 mm long post abscission, elevating the placental mass 2 mm above the persistent basal half of the fruit, exposing the seed-covered placenta which resembles a brush post-seed dispersal (Fig. 1F, 2C). *Seeds* not seen (dispersed, leaving only the filamentous 0.5 mm long funicles (Fig. 1F).

### DISTRIBUTION

Republic of Guinea, southern Simandou range.

### HABITAT AND ECOLOGY

Both sites for the new *Afrothismia* were in damp, black, humic soils in deeply shaded understorey of submontane forest adjacent to permanent or near-permanent streams, in association with other species of achlorophyllous mycotrophic plants, principally *Gymnosiphon fonensis*, but at the Boyboyba site, several other species also (see Discussion). Elevational range. c. 800 m.

### PHENOLOGY

In early October, at both sites, plants of the new species had open flowers, and flowers in bud, as well as both developing and old (post seed-dispersal) fruits. Plants are likely above ground July to October.

### CONSERVATION STATUS

Despite the fact that the two sites of *Afrothismia fonensis* are found in or immediately adjacent to the Pic de Fon Forêt Classée (PDF FC) of the southern Simandou range, a Tropical Important Plant Area (Couch et al. 2019, following the standard of Darbyshire et al. 2017), direct and indirect anthropogenic pressures represent major threats and the range of the species is taken as a single threat-based location in the sense of IUCN (2012). The PDF FC hosts a major open pit iron ore mine, the first ore being exported in late 2025. Both sites are on the footprint of the project. While the pit is not expected to directly impact sites for this species, there are risks that despite best efforts to protect threatened species, there will be negative impacts linked with the activities and infrastructure associated with the expected ore extraction, such as alteration of hydrology from de-watering of the adjacent Oueleba iron-ore body that is now being mined, and construction of new roads.

There has already been a slight decline in quality of habitat e.g. due to interventions for environmental surveys such as creation of paths (Cheek et al. 2024b). Of more urgent and more major concern are threats from foraging by red river hogs (damage seen at the Boyboyba site in Oct. 2025) and trampling of streamsides by zebu cattle herds that use forest streams as dry season drinking troughs. Both animal species are protected and present in ever-increasing numbers in the PDF FC (pers. obs. 2025).

The two forest sites for the species are separated north to south by c. 5.8 km of unsuitable savanna woodland habitat as measured on Google Earth. The largest number of individuals was recorded at the tunnel forest, the most northern site, where the area of the gallery forest canopy is about 0.005 km^2^. Here about six separate clusters of flowering and fruiting stems were recorded all within 1-2 m of the stream along a band about 10 m long. Each cluster was 10-20 cm diam. and consisted of 2 to 5 flowering or fruiting stems, usually with exposed bulbil clusters. It was assumed that each cluster represented a single individual that had spread over the years. Due to the rarity of the plants no excavations were made and every attempt was made to avoid compacting the soil in their vicinity. At the southern site, the Boyboyba forest, measures c.0.4 km^2^ in area. Here two separate plant clusters were seen, one at the foot of a tree in an area with a film of surface water, the other 4-5 metres distant at the edge of a stream. These two clusters were within the previously known ‘saprophyte’ plot of c. 30 m × 10 m described in Cheek et al. (2024b). Although the two forest patches are each far less than 1 km^2^ in area, in calculating the area of occupancy (AOO) we are required to use the IUCN (2012) mandatory grid cell-size of 4 km^2^, therefore AOO for *Afrothismia fonensis* is calculated as 8 km^2^. We follow the IUCN rule to estimate the Extent of Occurrence as the same. However, the actual area occupied by the species is likely to be in the order of 0.01 Ha. The ‘saprophyte’ plot of c. 30 m × 10 m (see above) resulted from targeted searches at the Boyboyba Forest for achlorophyllous mycotrophs over several days in 2008 and 2022 by specialists (including the first two authors) for species including *Gymnosiphon fonensis* which species was only found in this plot while other species of achlorophyllous mycotroph were found in multiple spots over several hectares within the forest.

We estimate that less than 50 mature (flowering-sized) individuals are known, qualifying the species as Critically Endangered under Criterion D 1.

Since there have been past, present and future threats, there is a single threat-based location, and EOO meets the threshold for Critically Endangered under criterion Criterion B, we here assess *Afrothismia fonensis* as Critically Endangered, CR B1ab(iii)+2 ab(iii).

It is to be hoped that this species will be searched for and found at other locations which would allow a lower extinction risk assessment than that made here. However, previous surveys in 2008 outside Simandou in the correct season, albeit targeted at another genus of achlorophyllous mycotroph, *Gymnosiphon*, failed to find *Afrothismia*. Yet sites such as Mt Bero and Mts Ziama where this species might be found have not yet been thoroughly surveyed at the correct season for achlorophyllous mycotrophs by specialists in this group. *Afrothismia* may yet be found at such sites, even though typically species of the genus have very small ranges and are often single-site endemics.

We recommend that a management plan is developed to ensure the survival of this species and is implemented following the protocols of Couch *et al*. (2022). This should include a public sensitisation and education programme, addressing the threats of red river hogs and cattle herds, ideally through constructing animal-proof fences around the plants. We also recommend annual monitoring of *Afrothismia fonensis* as well as the Critically Endangered *Gymnosiphon fonensis* (and also the Endangered *G. samouritoureanus*) populations to determine trends in survivorship and threats, potentially increased guarding of the forest habitat, and seedbanking. Cultivation and translocation of achlorophyllous mycotrophic flowering species such as *Afrothismia fonensis* has not yet been achieved but might be possible if sufficient resources and time can be allocated for researching this, for which the first step should be to determine the symbiotic mycorrhizal fungi and autotrophic partner species of the fungi upon which the plant depends. Without this information, planning for the translocation of any achlorophyllous mycotroph has a low chance of success.

### NOTES

It is difficult to be certain of the closest extant relatives of *Afrothismia fonensis* without molecular analysis or more abundant material for comparative purposes. Superficially, the species resembles in the small size and bicolored perianth tube *Afrothismia pusilla* Sainge & Kenfack of Mt Kala, near Yaoundé, Cameroon. However, the last species has a straight, erect rather than a zigzag perianth tube axis, and bears multicellular extensions from the tube not seen in *A. fonensis* (see Sainge et al. 2013). The new species also has similarities with its closest geographic neighbour, *A. hydra*. The two species are of similar size and colour, however *A. hydra* Sainge & T. Franke has a vertically oriented mouth, monomorphic perianth lobes and the perianth tube ornamentation is tuberculate on the dorsal surface (not horizontally oriented, dimorphic and ornamentation of longitudinal ridges on ventral surfaces and flanks). In overall architecture *A. fonensis* more closely resembles *A. mhoroana* Cheek of the Eastern Arc Mountains of Tanzania. However, the last species is coloured translucent-white & yellow, and the perianth tube is tuberculate (vs white and dark red, and with longitudinal ridges) (Cheek & Jannerup 2006).

## Discussion

*Afrothismia fonensis* is currently both the most northerly and westerly occurring species of the genus and family. This status was formerly held by *Afrothismia hydra* Sainge & T.Franke which is geographically the closest species to *A. fonensis* at its location in the Akure Forest, Oponomu, western Nigeria, c. 1550 km to the ESE of the Simandou location of *A. fonensis* and lying about a degree further south at 7º N.

### The Placentophore

In *Afrothismia fonensis* the placental column elongates from 0.25-0.5 mm long in the ovary to 4-5 mm long post-abscission, a 10 to 20 fold increase in length (see description above). This feature is a unique character of the Afrothismiaceae (Cheek et al. 2023) and is not known otherwise in Dioscoreales. The structure was first investigated and highlighted by Imhof & Sainge (2008) who termed it a ‘placentophore’ and speculated that it is an adaptation to favour seed dispersal. While in some species of *Afrothismia* (e.g. Cheek 2009) the placenta is only just raised beyond the fruit, in *A. fonensis* it is far exerted, rivalling the likely maximal extent seen in the genus, which seems to be *A. hydra*.

### Intertepal lobes

In several species of *Afrothismia*, e.g. *A. baerae* Cheek (Cheek 2004) each of the tepals has a hastate base, with two short flanking lobes that point in the opposite direction from the main part of the tepal lobe. Uniquely in the genus for the present, in *Afrothismia fonensis*, the flanking lobes at the base of each tepal appear to have fused with those of the adjacent tepals, making a single structure, here termed the intertepal lobe. These resemble the tepals except in length, texture and directionality.

### New discoveries of Central African plant genera in the Guinea Highlands of West Africa

The discovery of *Afrothismia fonensis*, in Guinea, far to the West of the previously known range of the genus (which is centred in Cameroon, Cheek et al. 2023a) follows the pattern of other recent discoveries. *Keita* (previously *Anacolosa*) *deniseae* (Olacaceae) a liana was also recently discovered from the submontane forest of the Simandou Range, (Cheek et al. 2024c) while its long-known sister species *K. uncifera* is centred in Gabon and adjacent countries. Similarly, the genus of small trees, *Ternstroemia* (Ternstroemiaceae/Pentaphylacaceae) was only known in Africa from as far west as Nigeria and Cameroon (Cheek et al. 2017), until *T. guineensis* Cheek was discovered in Kounounkan (Cheek et al. 2019). Likewise, the genus *Saxicolella* (Podostemaceae) was confined to Central Africa also, until two new species to science were found in Guinea, including *S. deniseae* Cheek (Cheek et al. 2022). The tree genus *Talbotiella* (Leguminosae) is also centred in Cameroon, with species in adjoining Nigeria and Gabon, and an isolated westernmost outlier in Ghana (Mackinder et al. 2010) until a new species was discovered inland of Conakry (van der Burgt et al. 2018). Less dramatically, the African genus *Virectaria*, with eight species, is centred in Cameroon, previously with only one very widespread species extending westwards to Guinea, while recently two new endemic species of the genus have been found in Kounounkan of Guinea (Simbiano et al. 2024 and Couch et al. 2019). Such discoveries illustrate how scientific knowledge of species in Guinea is far from complete, and how recent advances have contributed to the increase of documented plant species in Guinea from about 3,000 to nearly 4,000 species (Gosline et al. 2023a; 2023b).

### Boyboyba Forest: West African centre of diversity for mycoheterotrophic plants

The Boyboyba forest is well known as having the highest species diversity recorded in West Africa for achlorophllyous mycotrophic species with five species present (Cheek & van der Burgt 2010) to which we now add a sixth, *Afrothismia fonensis*. The five species previously recorded are: *Campylosiphon* (formerly *Burmannia*) *congestus* (C.H.Wright) Maas (Burmanniaceae), *Sebaea oligantha* (Gilg) Schinz (Gentianaceae), *Gymnosiphon longistylus* (Benth.) Hutch., *G. samouritoureanus* and *G. fonensis* (Burmanniaceae). All five species grow together with *Afrothismia fonensis* in the single ‘saprophyte’ plot of c. 30 m × 10 m in Boyboyba forest in which *Gymnosiphon fonensis* is also found.

The Boyboyba forest also hosts the largest known global populations of the Critically Endangered *Keetia futa* Cheek (Cheek et al. 2018) and of the Endangered *Keita deniseae* Cheek (Cheek et al. 2024c).

## Acknowledgements

The specimens and observations cited in this paper were collected with the support of Simfer S.A. under the framework of collaboration between RBG, Kew and under the Memorandum of Collaboration between RBG, Kew and Université Gamal Abdel Nasser-Herbier National de Guinée (MINRESI), i.e. UGANC-HNG. We especially thank Simfer Environment staff David Hamilton, Harry Nevard, Sekou Soumaoro and Thomas Williams for their support of the fieldwork. The authors thank their colleagues at UGAN-C-HNG for support.

We thank two anonymous reviewers for constructive comments on an earlier version of this manuscript

The authors declare they have no conflict of interest.

